# Brain pericytes derived from human pluripotent stem cells retain vascular and phagocytic functions under hypoxia

**DOI:** 10.1101/2025.04.10.648232

**Authors:** Youbin Kim, Mingzi Zhang, Allison Bosworth, Julia TCW, Lina R. Nih, Kassandra Kisler, Abhay P Sagare, Ruslan Rust

## Abstract

The integrity and function of the blood-brain barrier (BBB) are largely regulated by pericytes. Pericyte deficiency leads to BBB breakdown and neurological dysfunction in major neurological disorders including stroke and Alzheimer’s disease (AD). Transplantation of pericytes derived from induced pluripotent stem cells (iPSC-PC) has been shown to restore the BBB and improve functional recovery in mouse models of stroke and pericyte deficiency. However, the molecular profile and functional properties of iPSC-PC under hypoxic conditions, similar to those found in ischemic and neurodegenerative diseases remain largely unexplored. Here, we demonstrate that iPSC-PC under severe hypoxia retain essential functional properties, including key molecular markers, proliferation rates, and the ability to migrate to host brain vessels via function-associated PDGFRB-PDGF-BB signaling. Additionally, we show that iPSC-PC exhibit similar clearance of amyloid beta (Aβ) neurotoxins from AD mouse brain sections under both normoxic and hypoxic conditions. These findings suggest that iPSC-PC functions are largely resilient to hypoxia, highlighting their potential as a promising cell source for treating ischemic and neurodegenerative disorders.

## Introduction

Pericytes are mural cells that wrap around the endothelium, playing an important role in maintaining blood-brain barrier (BBB) integrity, regulating blood flow, promoting vessel maturation, and clearing toxic proteins in the brain.^1–8^ Loss of pericytes leads to BBB breakdown, which is associated with neuronal dysfunction and poor outcomes in major neurological disorders, including stroke and Alzheimer’s disease in both mice and humans. ^9–12^

Recently, protocols have been developed to generate pericytes from induced pluripotent stem cells (iPSC-PC).^13–15^ Transplantation of iPSC-PC has shown to enhance BBB integrity in mouse models of ischemic stroke^16^ and pericyte-deficient mice^15^. Furthermore, iPSC-PC exhibit a transcriptomic and proteomic signature similar to primary human brain pericytes.^13–15^

Despite these advances, molecular profiling of iPSC-PC has largely been explored under normoxic conditions, even though cell therapy applications often involve reduced cerebral blood flow causing hypoxic environments, such as those seen after ischemic stroke^17^ or in Alzheimer’s disease (AD)^18^. It remains unclear how hypoxia influences the molecular profile and functional properties of iPSC-PC.

In this study, we demonstrate that iPSC-PC retain the expression of canonical markers and show similar proliferation rates under severe hypoxia *in vitro*. Transcriptomic profiling revealed that while hypoxia-related pathways were upregulated, key functional components of iPSC-PC, including tight junction and adhesion junction molecules, remained unaffected by hypoxia. Functionally, hypoxic iPSC-PC retained the ability to migrate towards host brain capillaries, extend processes, and form hybrid human-mouse microvessels with functional associations. Additionally, normoxic and hypoxic iPSC-PC exhibited comparable abilities to phagocytose amyloid beta (Aβ) neurotoxins from AD mouse brain sections.

## Results

### Hypoxia does not impact the growth or canonical marker expression in iPSC-PC

We generated pericyte-like cells from IPSCs via Neural Crest Cell (NCC) intermediates as previously described^15^, and exposed iPSC-PC to hypoxia for one week (iPSC-PC_HYP_). We used a control group of iPSC-PC under normoxic conditions (iPSC-PC_NORM_). We confirmed that O_2_ levels were stably reduced to 1% oxygen in the hypoxic group throughout the experiment (**Fig 1B**). Proliferation rates of iPSC-PC_HYP_ were similar to those of control iPSC-PC_NORM_ at 3, 5, and 7 days after hypoxia induction (**Fig 1C**). We observed no significant differences in the percentage of total cells expressing the canonical pericyte markers NG2 (iPSC-PC_HYP_: 83.7%; iPSC-PC_NORM_: 79.4%, p > 0.05) and PDGFRB (iPSC-PC_HYP_: 82.1%; iPSC-PC_NORM_: 85.7%, p > 0.05) in fluorescence immunostaining (**Fig. 1 E, F**). Additionally, the fluorescence signal intensity of NG2 and PDGFRB expressing cells was comparable between both groups (all p > 0.05) (**Fig. 1E, G**).

**Fig. 1.**
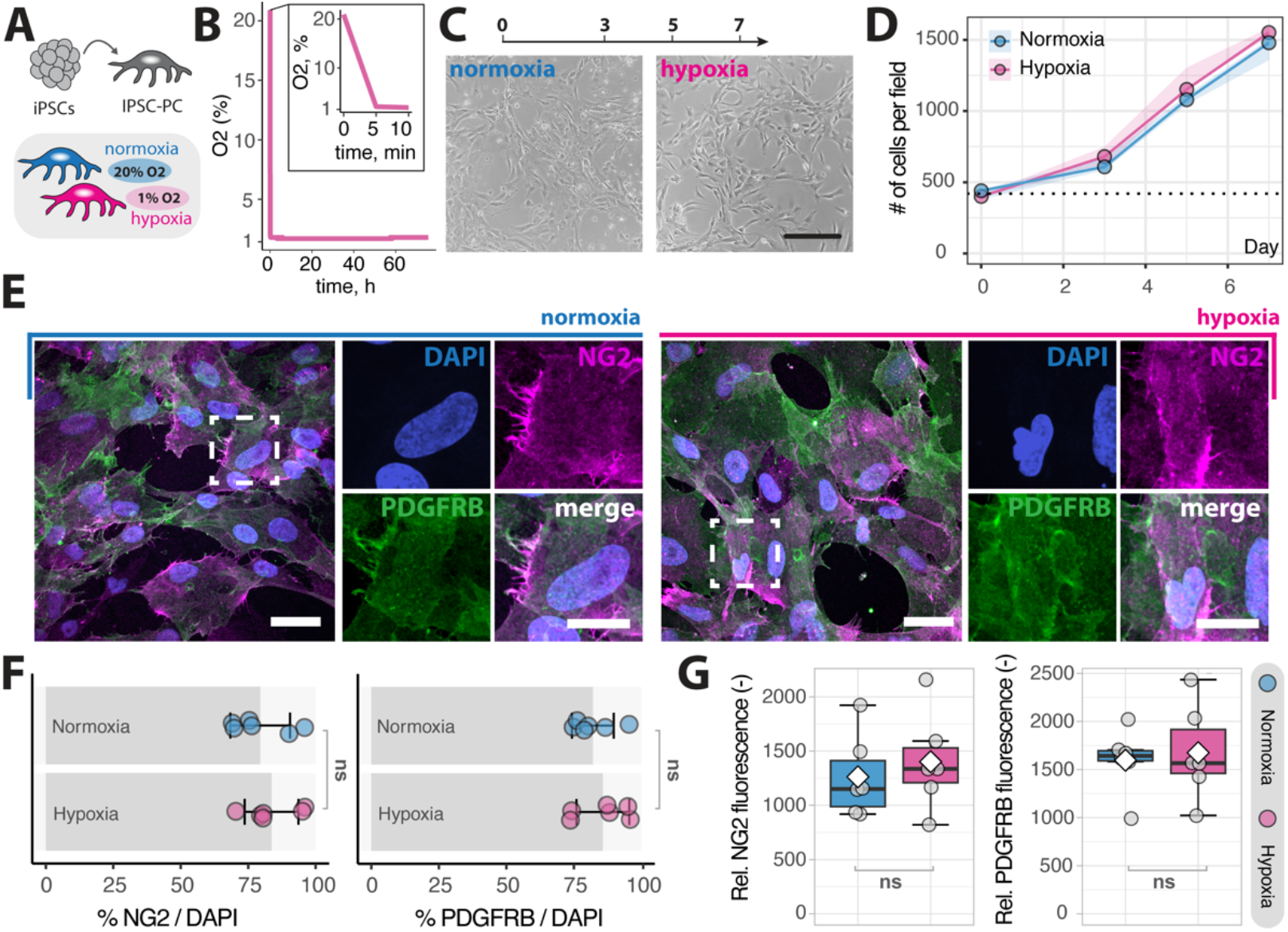
Growth and canonical marker expression of iPSC-PC under hypoxia. (A) Schematic overview of the experimental setup. (B) Oxygen concentration in the hypoxia chamber over 72 h. (C) Representative images of iPSC-PC at 7 days after hypoxia induction and normoxic iPSC-PC control. (D) Proliferation of iPSC-PC at 0, 3, 5, and 7 days after hypoxia induction. Scale bar: 50μm, N = 3. (E) Fluorescence images of iPSC-PCs stained for the canonical pericyte markers NG2 (magenta) and PDGFRB (green), counterstained with DAPI (blue). Scale bar: 10μm. (F) Quantification of the percentage of cells expressing NG2 and PDGFRB, N = 6. (G) Relative fluorescence intensity of NG2 and PDGFRB under normoxia and hypoxia, N = 6. Each dot represents an independent iPSC-PC culture. In line plots (D), error bars represent the standard error of the mean (SEM). In box plots (F,G), the box represents the interquartile range, the horizontal line indicates the median, the white diamond represents mean, and whiskers show the minimum and maximum values. Statistical analysis was performed using a repeated measures mixed model for (D) and an unpaired t-test for (F,G). ns = non-significant, p > 0.05.

These data suggest that hypoxia does not affect growth or canonical marker expression in iPSC-PC.

### The transcriptomic profile of iPSC-PC shows only minimal responses to hypoxia

To investigate the transcriptomic signature of iPSC-PC under hypoxia, we performed RNA-seq after 1 week of hypoxia. Overall, only 4.4% of all detected genes were differentially expressed genes (DEGs), with 1.7% upregulated and 2.7% downregulated in response to hypoxia (**Fig. 2A, B**). As expected, genes and pathways associated with hypoxia were upregulated, including gene ontology terms “response to decreased oxygen levels”, “response to oxygen levels” and “response to hypoxia” in iPSC-PC_HYP_ compared to IPSC-PC_NORM_ (**Fig. 2C**).

**Fig. 2.**
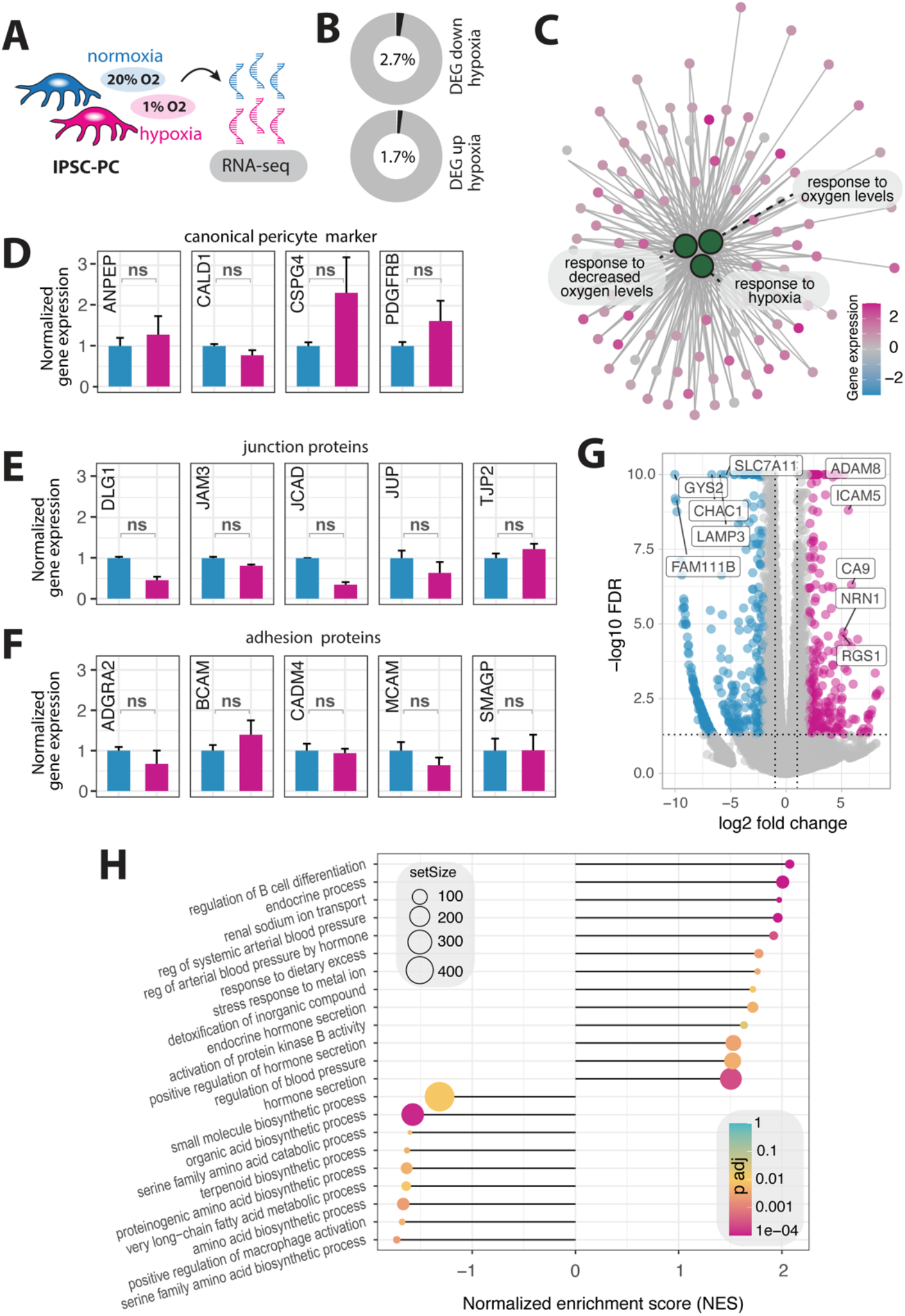
Transcriptomic analysis of iPSC-PCs under hypoxia. (A) Experimental setup for RNA sequencing after one week of hypoxia. (B) Percentage of differentially expressed genes (DEGs) in iPSC-PC_HYP_ compared to iPSC-PC_NORM_. (C) Gene interaction network highlighting hypoxia-responsive genes (small dots) within enriched pathways related to hypoxia (large green dots) upregulated in iPSC-PC_HYP_. (D) Normalized expression levels of canonical pericyte markers in iPSC-PC_HYP_ and iPSC-PC_NORM_. (E) Expression levels of key junction proteins in iPSC-PC_HYP_ and iPSC-PC_NORM_. (F) Expression levels of adhesion in iPSC-PC_HYP_ and iPSC-PC_NORM_. (G) Volcano plot showing differentially expressed genes (DEG), with selected DEG labeled (magenta, upregulated; blue, downregulated). (H) Gene set enrichment analysis (GSEA) showing most upregulated and downregulated in iPSC-PC_HYP_ and iPSC-PC_NORM_. Data represent N = 6 independent cultures. Genes with FDR < 0.05 and |log2 FC| > 1.5 were considered significantly differentially expressed, ns = non-significant.

Next, we explored whether genes encoding for canonical pericyte markers including PDGFRβ (*PDGFRB*), Caldesmon (*CALD1*), NG2 (*CSPG4*), CD13 (*ANPEP*), and Vitronectin (*VTN*) changed in response to hypoxia. Consistent with the histology data (**Fig. 1**), we did not detect significant gene expression changes in canonical pericyte markers between iPSC-PC_HYP_ and iPSC-PC_NORM_ (all FDR > 0.05, **Fig. 2D**). We then examined whether hypoxia influenced the expression of genes encoding for major functional components of iPSC-PC including cell adhesion and tight junction proteins (**Fig. 2E, F)**. We found no significant changes in the expression of genes encoding for major junction proteins (*DLG1, JAM3, JCAD, JUP, TJP2*, all FDR > 0.05) and adhesion junction proteins (*ADGRA2, BCAM, CADM4, MCAM, SMAGP*, all p > 0.05) (**Fig. 2E, F)**. Among the genes most significantly upregulated upon hypoxia, we detected *ICAM5* (log_2_FC = 1.92, FDR < 0.001), *HIF3A* (log_2_FC = 5.60, FDR < 0.001), *CA9* (log_2_FC = 4.35, FDR = 0.001), and *TEK* (log_2_FC = 2.08, FDR < 0.001), which are known regulators of vascular adaptation to hypoxia (**Fig 2G**). Conversely, genes downregulated in response to hypoxia included *GYS2* (log_2_FC = -6.69, FDR < 0.001), *CHAC1* (log_2_FC = -5.87, FDR < 0.001), *LAMP3* (log_2_FC = -5.87, FDR < 0.001), and *CD36* (log_2_FC = -6.90, FDR = 3.08e-02), which are associated with reduced glucose and lipid metabolic processes (**Fig 2G**).

Given that hypoxia is known to influence various cellular processes beyond oxygen response, we conducted a gene set enrichment analysis (GSEA) to identify additional pathways affected by hypoxia. Primary pathways that were upregulated in response to hypoxia, aside from those related to hypoxia, included endocrine and hormonal regulation, immune responses, and vascular regulators (**Fig. 2H**). In contrast, downregulated pathways were associated with biosynthetic processes including amino acid and lipid metabolism (**Fig. 2H**), which is consistent with the observed DEGs.

Collectively, these data suggest that while hypoxia triggers expected oxygen-related responses and pathways associated with metabolism, it does not significantly impact the gene expression of canonical pericyte markers or key structural and functional components.

### iPSC-PC functionally home to the mouse brain vasculature under hypoxia

The ability of iPSC-PC to wrap around brain capillaries is a prerequisite for pericyte-based therapies to restore the BBB integrity. To evaluate whether iPSC-PC retain their ability to functionally home to the mouse brain capillaries and form hybrid-microvessels under hypoxic conditions, we used acute living brain slices from pericyte deficient *Pdgfrb*^F7/F7^ mice and incubated them for 24 h with iPSC-PC under three conditions: (1) iPSC-PC_NORM_ with normoxic brain slices (iPSC-PC_NORM_ + BRAIN_NORM_) (2) iPSC-PC_HYP_ with normoxic brain slices (iPSC-PC_HYP_ + BRAIN_NORM_), and (3) iPSC-PC_HYP_ with hypoxic brain slices (iPSC-PC_HYP_ + BRAIN_HYP_) to mimic the hypoxic host environment (**Fig. 3A, B**).

**Fig. 3:**
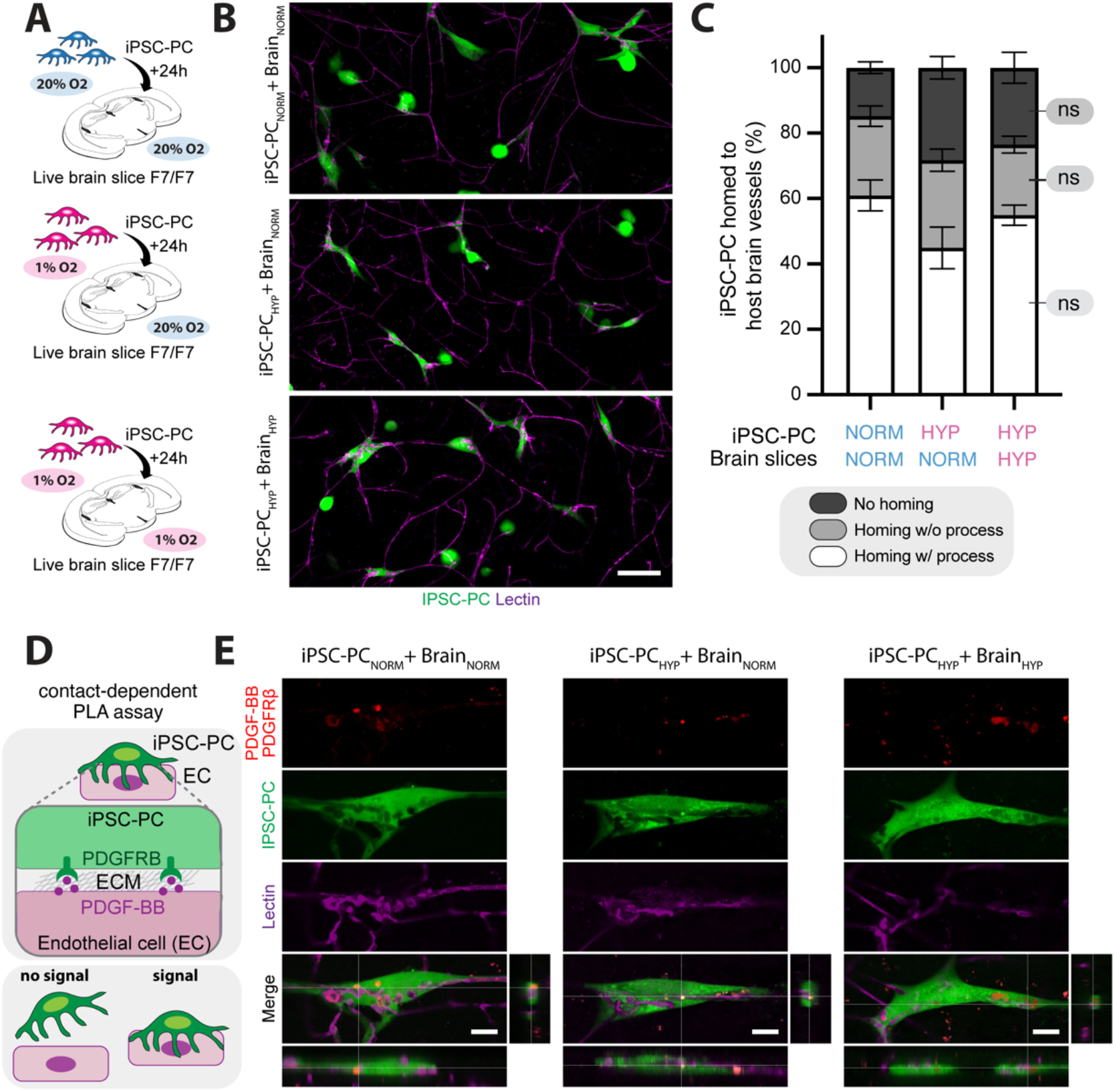
IPSC-PC home to brain capillaries and form functional hybrid microvessels under normoxia and hypoxia. (A) Schematic overview of experimental setup. Acute brain slices from pericyte-deficient *Pdgfrb*_F7/F7_ mice were incubated with iPSC-PC under three conditions: : 1) PSC-PC_NORM_ + BRAIN_NORM_, (2) PSC-PC_HYP_ + BRAIN_NORM_, and (3) PSC-PC_HYP_ + BRAIN_HYP_. (B) Representative confocal images showing iPSC-PC (green, CellTracker) interacting with host brain capillaries (magenta, Lectin). Scale bar: 20 μm. (C) Quantification of IPSC-PC homing and process formation with the host brain endothelium across conditions. (D) Schematic of the proximity ligation assay (PLA) used to assess pericyte-endothelial cell contact of PDGFRB (pericytes) and PDGF-BB (endothelium). (E) Representative PLA images showing PDGFRB-PDGF-BB colocalization (red) at the pericyte-endothelial interface under all three conditions, confirming functional interaction. Scale bar: 5 μm. Data represent N = 4 independent cell cultures for each condition. Statistical significance was determined by one-way ANOVA followed by Tukey’s post hoc test, ns = non-significant.

We first quantified the percentage of iPSC-PC that homed to the host brain endothelium, distinguishing between those cells that formed processes along the vessels and those without processes. Across all groups, we found that a similar proportion of iPSC-PC homed with processes (iPSC-PC_NORM_ + BRAIN_NORM_: 60.9 ± 4.7%, iPSC-PC_HYP_ + BRAIN_NORM_: 44.9 ± 6.3%, iPSC-PC_HYP_ + BRAIN_HYP_: 54.9 ± 3.1%, p > 0.05) and homed without forming processes (iPSC-PC_NORM_ + BRAIN_NORM_: 24.3 ± 3.1%, iPSC-PC_HYP_ + BRAIN_NORM_: 26.8 ± 3.4%, iPSC-PC_HYP_ + BRAIN_HYP_: 21.5 ± 2.6%, p > 0.05). Only a minor fraction of iPSC-PC failed to home in all experimental groups (iPSC-PC_NORM_ + BRAIN_NORM_: 14.8 ± 1.8%, iPSC-PC_HYP_ + BRAIN_NORM_: 28.3 ± 3.5%, iPSC-PC_HYP_ + BRAIN_HYP_: 23.5 ± 4.7%, p > 0.05) (**Fig. 3C**).

Next, we aimed to investigate whether the iPSC-PC can form functional interactions with the host endothelium of Pdgfrb_F7/F7_ mice. We performed a proximity ligation assay (PLA) targeting known pericyte-endothelial cell interaction molecules, PDGFRB and PDGF-BB^19–21^ (**Fig 3D**). PLA signal of PDGFRB/PDGF-BB was detectable in all three experimental groups at the pericyte-endothelial interface (PDGFRB^+^ Lectin^+^), suggesting the formation of functional pericyte-endothelial contacts in all three experimental conditions (**Fig. 3E**).

### IPSC-PC phagocytic properties of Aβ neurotoxins remain unchanged under hypoxia

Previous studies suggest that pericytes can phagocytose Aβ *in vitro*.^15,22,23^ Therefore, we aimed to test whether iPSC-PC have comparable Aβ clearance properties under normoxic and hypoxic conditions. We used consecutive frozen brain sections from 10-month old 5xFAD mice, which are known to have significant Aβ pathology at this age^24,25^, and incubated these sections with iPSC-PC_NORM_ and iPSC-PC_HYP_ to these sections for 24 hours. Brain sections from 5xFAD mice without cells served as a negative control (**Fig. 4A**). After confirming the presence of iPSC-PC on the brain sections after 24 hours, we quantified the Aβ load in the cortex and hippocampus (**Fig. 4B**). We observed a similar reduction in Aβ levels, which decreased from 32.2 ± 1.5% in the control group to 21.7 ± 0.5% in iPSC-PC_NORM_, (p < 0.01) and to 23.9 ± 1.7% in iPSC-PC_HYP_, (p < 0.05) in the cortex (**Fig. 4C**). A similar trend was observed in the hippocampus, with Aβ levels reduced from 24.3 ± 1.3% in the control to 20.1 ± 0.7% (iPSC-PC_HYP_, p = 0.2, Fig. 4C) after cell incubations. Notably, no significant differences in Aβ uptake were observed between iPSC-PC under normoxic and hypoxic conditions, suggesting similar phagocytic properties in both environments (all p > 0.05, **Fig. 4C**).

**Fig. 4:**
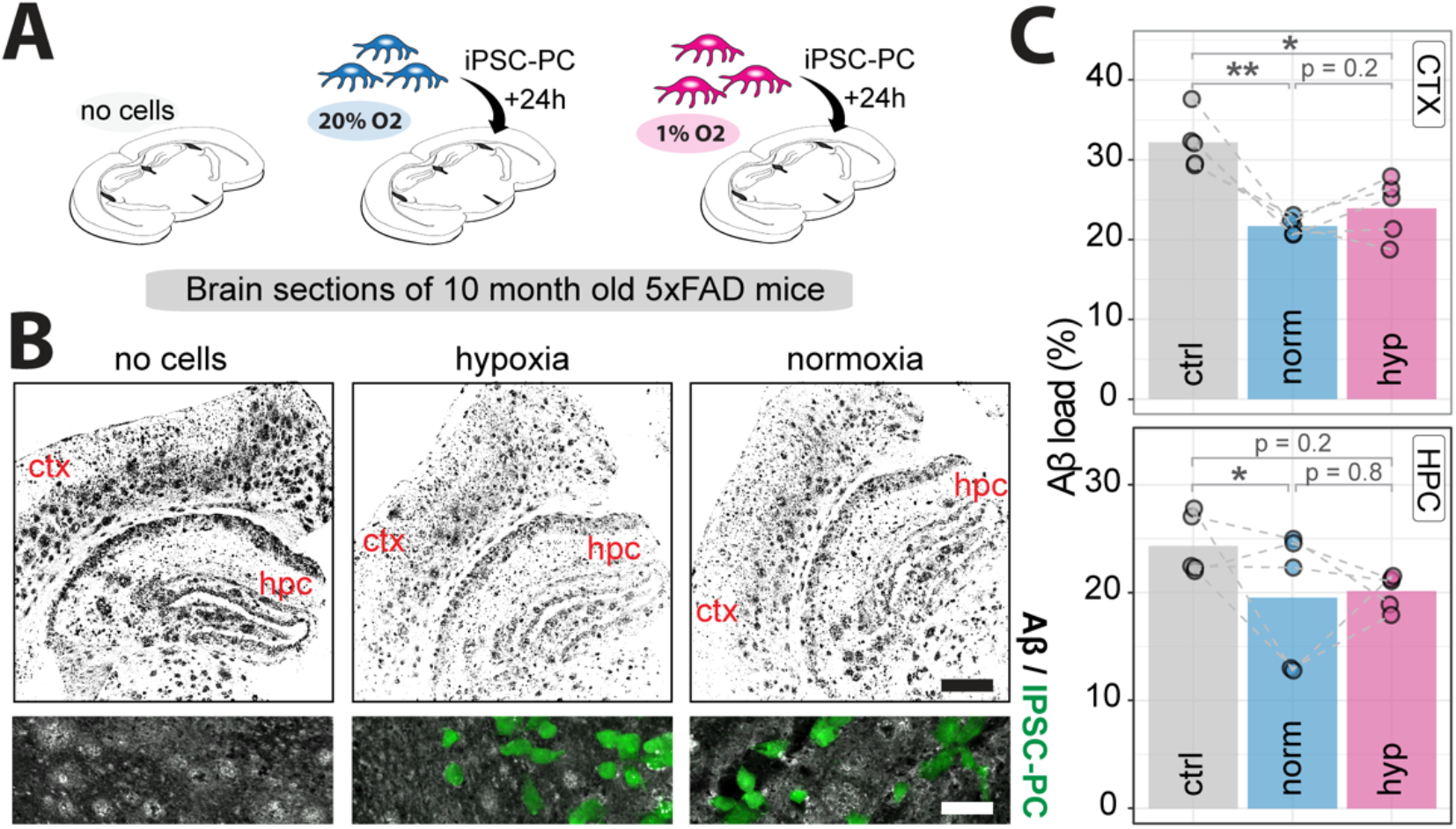
IPSC-PC phagocytose Aβ from brain sections of aged 5xFAD mice under normoxia and hypoxia. (A) Schematic overview of experimental setup. Frozen brain sections from 10-month-old 5xFAD mice were incubated with iPSC-PC under three conditions: (1) no cells, (2) iPSC-PC_NORM_, and (3) iPSC-PC_HYP_. (B) Representative confocal images showing Aβ (black). Magnification view within the section showing iPSC-PC (green, CellTracker) and Aβ (gray). Scale bar: overview: 200 μm, zoom in: 50 μm (C) Quantification of Aβ load (%) in the cortex (CTX) and hippocampus (HPC). Each dot represents a individual brain section, with dotted lines connecting paired consecutive sections from the same brain region across conditions. Data represent N = 5 independent cell cultures for each condition. Bars represent mean, each dot represents iPSC-PC added to a different brain slice. Dashed lines indicate consecutive brain sections. Asterisks indicate significance: *p < 0.05, **p < 0.01 using paired t-test with correction for multiple testing.

## Discussion

In this study, we show that iPSC-PC retain their major molecular and functional properties under one week of severe hypoxia. We identified similar growth rates, expression of canonical pericyte markers and junctional and adhesion proteins were unaffected by hypoxia. Additionally, IPSC-PC can home to pericyte-deficient host vessels in a hypoxic environment and have similar phagocytic properties of Aβ to normoxic iPSC-PC.

Traditionally, brain pericytes have been considered to be highly susceptible to hypoxia as they are often the first non-neural cell types lost after severe hypoxia e.g., after experimental stroke^26–29^ or pathological CBF changes in dementias^30–32^ contributing to BBB disruption and subsequent neurodegeneration^1^. Therefore, transplantation of pericytes is an interesting therapeutic target to restore the BBB, however it was uncertain whether iPSC-PC can retain their functional properties in the hypoxic microenvironment. While some recent studies showed that transplantation of pericytes can enhance the BBB integrity in mouse models of ischemic stroke^16^ and in pericyte-deficient mice^15^, most studies used mesoderm-derived pericytes. However, in major neurological disorders including stroke and AD, pericyte degeneration is observed in the cortex and hippocampus, which primarily affects forebrain NCC-derived pericytes, but not mesodermal pericytes.^33^ Our data now shows that NCC-derived iPSC-PC, which have been previously shown to closely resemble NCC-derived primary forebrain pericytes^15^, can also sustain a hypoxic environment.

We observed in hypoxic iPSC-PC energy metabolic reprogramming, which has been previously observed in vascular cells, favoring e.g. glycolysis over oxidative phosphorylation.^34,35^ Additionally, upregulation of *ICAM5, HIF3A, CA9*, and *TEK* in hypoxia-exposed pericytes could suggest an adaptive mechanism to maintain vascular support.^36,37^ Furthermore, hypoxic iPSC-PC maintained expression of the important pericyte-endothelial communication signaling of PDGFRB and PDGF-BB, which has been previously described to be important for maintaining the BBB-supporting functions of pericytes.^19–21^ While we identified the molecular signature of hypoxic iPSC-PC and their interaction with endothelial cells, we used a simplified *in vitro* model with reduced (1% O2) concentration and *ex vivo* brain slices, however *in vivo* validation should be performed in future studies to assess iPSC-PC physiological integration to the host vasculature and therapeutic efficacy. Additionally, while our study examined the response of iPSC-PC for one week after hypoxia, prolonged hypoxia may induce additional changes in iPSC-PC that need further exploration.

Previous work has shown that pericytes can phagocytose Aβ *in vitro*^15,22,23^, which is in accordance with our findings. We additionally show that iPSC-PC retain their Aβ phagocytic properties under hypoxia indicating that iPSC-PC could take up Aβ even in the presence of a low-oxygen environment in Alzheimer’s Disease. Future studies could confirm these findings in AD mouse models to assess iPSC-PC therapeutic potential *in vivo*. Potentially, the ability of iPSC-PC to improve the vascular and Aβ pathology in AD could also be tested in new AD mouse models with a stronger vascular phenotype.^38^

In conclusion, the present study provides support that iPSC-PC maintain their major functional properties under hypoxia. Therefore, iPSC-PC might be a suitable cell source for future brain transplants for neurological disorders that are associated with pericyte deficiency and hypoxia such as stroke, AD and other related disorders.

## Methods

### Mice

Platelet-derived growth factor receptor β mutant mice, Pdgfrb^F7/F7^, on 129S1/SvlmJ background were used for ex vivo and in vivo studies, as described below. Mice express PDGFRb with seven point mutations that disrupt signal transduction pathways; including residue 578 (Src), residue 715 (Grb2), residues 739 and 750 (PI3K), residue 770 (RasGAP), residue 1008 (SHP-2), by changing the tyrosine to phenylalanine, and residue 1020 (PLCγ), where tyrosine was mutated to isoleucine, as reported ^*39*^. Pdgfrb^F7/F7^ mice express mutant PDGFRβ exclusively in perivascular mural cells including pericytes, and not in neurons, astrocytes or brain endothelial cells ^*3,40*^. All procedures received approval from the Institutional Animal Care and Use Committee at the University of Southern California in accordance with U.S. National Institutes of Health guidelines and were conducted following the ARRIVE guidelines^41^.

#### Cell culture

Human iPSC cultures were generated from skin fibroblasts from as previously described ^42^. Human iPSC-PC were generated via NCC intermediates following previously established protocols^43,44^. Briefly, iPSC were cultured on growth factor reduced matrigel (Corning, 356230) in mTeSR Plus medium (StemCell Technologies, 100-0276). NCC differentiation was induced using STEMdiff™ Neural Crest Differentiation Kit (StemCell Technologies, 08610). Prior to differentiation, iPSC were washed with PBS, dissociated with Accutase (Thermo Fisher, A1110501) for 5 min, centrifuged at 200xg for 4 min, and counted using a Countess 3 (Thermo Fisher, AMQAX2000). IPSCs were then seeded at a density of 0.75-1×10^5^ cells/cm^2^ in mTeSR plus (StemCell Technologies, 1000276) supplemented with 10 mM Y-27632 (Tocris, 1254). Seeding densities were adjusted for each iPSC line to ensure 100% confluency at differentiation day 3-4. Media was changed daily using NCC media from the STEMdiff kit. After six days of differentiation, iPSC-NCC were dissociated using Accutase for 5 min and purified using EasySep™ Release Human PSC-derived NCC Positive Selection Kit (StemCell Technologies, 100-0047), following the manufacturer’s instructions.

NCC were further differentiated to iPSC-PC. IPSC-derived NCC were replated onto matrigel-coated 6-well plates at a 1:2 split ratio, and pericyte medium (ScienCell, 1201) was introduced the following day. Media were changed daily, and once cells reached 70-80% confluence, they were passaged using 0.05% Trypsin/EDTA (Thermo Fisher, 25300054) and replated onto poly-L-lysine (PLL)-coated dishes. After 14 days of differentiation, iPSC-PC were transferred onto glass coverslips for subsequent imaging.

For hypoxia induction, iPSC-PC from the same cell line were cultured under either normoxic (21% O_2_) or hypoxic (1% O_2_) conditions for up to 7 days using a Hypoxia Incubator Chamber (STEMCELL Technologies, 27310) according to the manufacturer’s instructions. After hypoxia treatment, cells were either fixed for immunocytochemical analysis, processed for RNA sequencing, or added to brain tissue for *ex vivo* assays.

#### Immunostaining

IPSC-PC were plated on glass coverslips and maintained in culture before fixation with 4% PFA for 10 minutes. Cells were then rinsed twice with sterile PBS, followed by three consecutive 5-minute washes in PBS. Blocking was performed for at least 1 hour in PBS containing 5% donkey serum and 0.5% Triton X-100. Cells were incubated overnight at 4°C with primary antibodies against PDGFRβ (R&D Systems, AF385, 1:100, Goat) and NG2 (Sigma, AB5320, 1:500, Rabbit) diluted in PBS supplemented with 5% donkey serum (PBSD). The following day, cells underwent five 8-minute washes in PBS before incubation with secondary antibodies (Invitrogen, A32794, Alexa Fluor Plus 555, 1:300, and Invitrogen, A32860, Alexa Fluor Plus 680, 1:300, Donkey) in PBSD for 1–2 hours at room temperature. After four additional 8-minute PBS washes, coverslips were mounted with DAPI-containing mounting medium (Southern Biotech, 0100-20) and imaged using a Nikon A1R confocal microscopy system with NIS-Elements software, using 10×, 20×, and 60× objectives.

#### Quantification of PDGFRB- and NG2-positive cells

All samples were stained and imaged using identical image acquisition settings; images were processed and analyzed using ImageJ (Fiji), similar to previous studies ^30^. Fluorescence signal from the marker of interest (PDGFRB, NG2) was thresholded using Otsu thresholding plugin ^45^ in each image. The thresholding was performed in the same manner for all samples. Analyze Particles function was used to determine the number of marker-positive cells. To calculate the percentage of positive cells, the number of marker-positive cells was normalized to the total DAPI-positive nuclei count for each image.

#### RNA sequencing and analysis

Total RNA from iPSC-PC was extracted using RNeasy RNA isolation kit (Qiagen, 73404), including DNase treatment to remove residual genomic DNA, according to the manufacturer’s instruction, and as previously described ^26,29,46–48^. All samples had an RIN value greater than 8.5. Library preparation, sequencing, read processing, alignment, and read counting were performed at the USC Norris Cancer Center Molecular Genomics Core. For library preparation, the TruSeq Stranded RNA kit (Illumina Inc.) was used following the manufacturer’s protocol. mRNA was purified via polyA selection, chemically fragmented, and transcribed into cDNA before adapter ligation. Initial data preprocessing, including quality control, adapter trimming, and filtering of low-quality reads, was performed using Galaxy open source platform. Reads were aligned to the human genome (GRCh38) using STAR aligner with default parameters. RNA sequencing data analysis and gene set enrichment analysis were conducted as previously described ^29,46,49^ and using standard guidelines with EdgeR^50^ and clusterProfiler 4.0^51^ with default parameters, applying the Benjamini-Hochberg (BH) method for multiple testing correction.

#### Ex vivo living brain slices preparation and analysis

We used acute living brain slice culture as a method to study IPSC-PC–vascular interactions in a physiologically relevant brain environment *ex vivo*. For this assay, iPSC-PC were labeled with CellTracker Green CMFDA (Invitrogen, C7025) following the manufacturer’s instructions, and cultured in either normoxic or hypoxic condition for one week before the brain slice preparation. The 8-month-old *Pdgfrb*^*F7*/*F7*^ mice were retroorbitally injected with 80 μL DyLight-649 labeled Lycopersicon esculentum lectin (Vector Laboratories, DL-1178-1)^3^ to identify blood vessels at least 15 min prior to euthanasia. Mice were deeply aestheticized with 5% Isoflurane, then, rapidly decapitated and brain extracted into ice-cold Hibernate A minus Phenol Red medium (Transnetyx Tissue, HAPR500). Slice preparation was performed on a Leica VT1000S vibratome. Coronal slices (250 μm) containing the hippocampus were then cut in ice-cold Hibernate A minus Phenol Red medium.

After sectioning, slices were maintained in homing medium containing DMEM (Gibco, 10569010), 5% FBS (Gibco, 16140-071), Pen-Strep (Gibco, 15140122), 2% platelet lysate (Stem Cell Technology, 200-0323), anti-oxidant (Sigma-Aldrich, A1345), GlutaMAX (Gibco, 35050061), Na-Pyruvate (Gibco, 11360070), CultureBoost (Cell Systems, 4CB-500), Pericyte Supplement (ScienCell, 1252). Subsequently, 50,000 iPSC-PC were added on top of each brain slice and allowed to integrate with the tissue in an incubator maintained at 37°C in either normoxic or hypoxic conditions for 24 h. Brain slices were fixed in 4% PFA for 30 min, mounted on slides, coverslipped with mounting media and imaged on a Nikon A1R confocal microscopy system with NIS-Elements software control. The iPSC-PC on brain slices were scored as a) being associated with a vessel and forming processes, b) associated with a vessel without processes or c) not associated with a vessel.

#### Proximity ligation assay (PLA)

The interactions of PDGF-BB and PDGFRβ in the brain slices were determined by proximity ligation assay using NaveniFlex Tissue GR Red (Navinci, 39220). After fixation and blocking, brain slices were incubated with rabbit anti-PDGF-BB IgG (Invitrogen, MA5-51346, 1:100) and goat anti-human PDGFRβ IgG (R&D Systems, BAF385, 1:50) overnight at 4ºC. Proximity ligation was then conducted following the manufacturer’s instructions. Proximity ligation results in a fluorescence signal only when the PLA agent detects the antibodies for PDGFRβ and PDGF-BB within close proximity, indicating likely colocalization of PDGFRβ and PDGF-BB.

#### Aβ *uptake assay from brain sections*

IPSC-PC labeled with CellTracker Green CMFDA (Invitrogen, C7025) and cultured in either normoxic or hypoxic conditions for 1 week before the brain slice preparation. Cryosections (20 μm thick) from 10-month-old *5xFAD* mouse brains were mounted on poly-L-lysine (PLL)-coated coverslips and stored at -80°C. Prior to use, sections were thawed, rinsed with PBS, and incubated in pericyte culture medium (ScienCell) for 2 hours at 37°C. IPSC-PC were seeded onto tissue sections at 100k cells/well, with ‘no-cell’ controls included. Cultures were maintained for 24 hours at 37°C. Sections were then fixed in 4% paraformaldehyde, permeabilized in PBS containing donkey serum and 0.075% Triton X-100, and stained with a pan-Aβ antibody (1:1200, Cell Signaling Technology) followed by secondary antibody incubation. Coverslips were mounted using DAPI-containing mounting medium and imaged using a confocal microscope at 4x magnification. For analysis, images were binarized using a customized threshold and percentage of Aβ signal were quantified in cortex and hippocampus using ImageJ (Fiji).

#### Statistics

Data are presented as mean ± standard error of the mean (SEM). All analyses were performed using GraphPad Prism 10 or R Studio 3.6.0. Normality of the data was tested using the Shapiro-Wilk test. For comparisons between two groups, an unpaired two-tailed Student’s t-test was used for independent samples, and a paired t-test was applied for repeated measures. For multiple group comparisons, a one-way analysis of variance (ANOVA) was used, followed by Tukey’s post hoc test for pairwise comparisons. For repeated measures across different groups, a repeated measures mixed model was applied. For quantification of gene expression changes in transcriptomic data, differential gene expression analysis was performed using EdgeR, and multiple testing correction was applied using the Benjamini-Hochberg (BH) method. Statistical significance was defined as *p < 0.05, **p < 0.01, and ***p < 0.001.

## Data availability

All data are available upon request. RNA-seq data will be made publicly available on NCBI GEO.

## Acknowledgements

-

## Supplementary material

-

## Contributions

RR contributed to overall project design, YK, MZ, AB, RR, AS performed the experiments. YK, MZ, AB, LRN, RR analyzed the data, JTCW generated iPSC lines, KK, AS, RR supervised the study. RR made figures. RR wrote the manuscript with input from all authors. All authors read and approved the final manuscript.

